# Sex-specific maladaptive responses to acute stress upon in utero THC exposure are mediated by dopamine

**DOI:** 10.1101/2024.09.17.613444

**Authors:** Serra Valeria, Traccis Francesco, Aroni Sonia, Vidal Palencia Laura, Concas Luca, Serra Marcello, Leone Roberta, Porcu Patrizia, Busquets Garcia Arnau, Frau Roberto, Melis Miriam

## Abstract

Cannabis remains by far the most consumed illicit drug in Europe. The availability of more potent cannabis has raised concerns regarding the enhanced health risks associated with its use, particularly among pregnant women. Growing evidence shows that cannabis use during pregnancy increases the risks of child psychopathology. We have previously shown that male rat offspring prenatally exposed to Δ^9^-tetrahydrocannabinol (THC), a rat model of prenatal cannabinoid exposure (PCE), display a hyperdopaminergic phenotype associated with a differential susceptibility to acute THC- and stress-mediated effects on sensorimotor gating functions. Here, we explore the contribution of the hypothalamic-pituitary-adrenal (HPA) axis, key regulator of body adaptive stress responses, to the detrimental effects of acute stress on ventral tegmental area (VTA) dopamine neurons and sensorimotor gating function of PCE rats. We report a sex-dependent compromised balance in mRNA levels of genes encoding mineralocorticoid and glucocorticoid receptors in the VTA, alongside with stress-induced pre-pulse inhibition (PPI) deficits. Notably, VTA dopamine neuronal activity is required for the manifestation of stress-dependent deterioration of PPI. Finally, pharmacological manipulations targeting glycogen-synthase-kinase-3-β signaling during postnatal development correct these stress-induced, sex-specific and dopamine-dependent deficits of PPI. Collectively, these results highlight the critical sex-dependent interplay between HPA axis and dopamine system in the regulation of sensorimotor gating functions in rats.

## 1. Introduction

Mental disorders affect 25% of the population worldwide and more than one-third of these are established by 14.5 years of age [1–3]. Stress is widely accepted as a detrimental factor for mental wellbeing, leading the World Health Organization to classify it as the “health epidemic of the 21st century” [4], with stress-related mental illnesses progressively increasing worldwide, particularly in youth [5]. However, it is still unclear why some individuals are more susceptible to developing stress-induced psychopathology, while others are more resilient and/or recover more rapidly [6, 7].

Many factors regulate our ability to cope with stress, including inter-individual differences due to genetic background, gender/sex, and early life adversities. Adaptive responses to stress can be modified by experience, which can increase the sensitivity of the hypothalamic-pituitary-adrenal (HPA) axis and dopamine system and produce inadequate stress reactions [8, 9]. When adverse experiences occur early in life, they might impact on developmental trajectory of different systems and circuitries with long-lasting and profound effects [7, 10, 11]. A functional interplay between the HPA axis and dopamine system plays an important role for adequate coping strategies in reaction to stressful conditions [8, 9, 12, 13]. Preclinical studies have demonstrated that acute stressors activate dopamine neurons [14] promoting long-lasting neuroplastic changes [15, 16] and leading to a gain-of-function of dopamine signaling that might alter the individual coping strategies to subsequent insults later in life [13]. Acute stress is also associated with mesolimbic dopamine release in humans [17]. Understanding how early life adversities contribute to both vulnerability to and resilience against acute stress and manifestation of psychopathology is crucial to implement early diagnosis and treatment, and to improve clinical outcome [7, 18].

Clinical evidence shows that among early life adversities prenatal cannabis exposure (PCE) is a predictive risk factor for the development of offspring psychopathology [19–26], and is associated with alterations in cortisol levels [27–29]. However, cannabis use during pregnancy has been on an alarming sharp rise [30, 31], with the greatest use during the first trimester [32]. Indeed, cannabis increasing legal availability is most likely leading to the common misconception that it is a safe natural remedy to be used even during pregnancy [33–35]. We have previously found that, in a rat model of PCE, male but not female subjects manifest an at-risk psychopathological endophenotype for a quantifiable trait of deficits in sensorimotor gating functions upon a single exposure to the cannabis main psychoactive ingredient, delta9-tetrahydrocannabinol (THC) [36]. This is associated with a sex-specific mesolimbic hyperdopaminergia [36, 37], a hallmark of disrupted sensorimotor gating functions [38–41]. We also found that pre-pulse inhibition (PPI) of the startle reflex [42, 43], a cross-species validated operational measure of sensorimotor gating functions, was impaired in PCE males upon either an otherwise ineffective dose of dopamine receptor agonist apomorphine or an acute stressor [44]. Of note, sex-specific failure in adopting coping strategies in response to an acute unescapable stressor were manifest as a consequence of PCE [37, 44].

In this study, we investigated how sex contributes to a PCE-induced vulnerability to the effects of acute stress. By using behavioral and molecular analyses, we found that PCE progeny exhibit male-specific reduced feedback sensitivity associated with exaggerated responses to an acute inescapable stressor. By using chemogenetic approaches, we showed the causal role of dopamine neuron activity of the ventral tegmental area (VTA) in the deteriorating effects of acute inescapable stressors on sensorimotor gating functions. Finally, such maladaptive stress responses in PCE males could be prevented by postnatal treatment with either pregnenolone or lithium, two inhibitors of glycogen synthase kinase-3 beta (GSK-3β) signaling pathway. Our data expand our understanding of the sex-specific impact of PCE on dopamine neurodevelopmental trajectories suggesting molecular targets for therapeutic intervention of offspring exposed to cannabis during pregnancy and suggest a causal link between a hyperdopaminergic profile and stress-induced deficits of PPI.

## 2. Materials and Methods

### 2.1 Subjects

All experimental procedures were performed in accordance with the European legislation EU Directive 2010/63 and were approved by the Animal Ethics Committees of the University of Cagliari and by the Italian Ministry of Health (Authorization n° 256/2020). We made all efforts to minimize pain and suffering and to reduce the number of animals.

### 2.2 Drugs and treatments

#### 2.2.1 Drugs and chemicals

THC resin was purchased from THC PHARM GmbH (Frankfurt, Germany) and dissolved in ethanol at 20 % final concentration. THC was diluted with sterile saline (0.9 % NaCl) containing 1-2 % Tween® 80. Clozapine-N-oxide dihydrochloride (CNO, Tocris Bioscience) was dissolved in sterile saline (0.9% NaCl). CNO was administered intra-peritoneally (i.p.) at 2 ml per kg body weight, 30 minutes prior to PPI test. Pregnenolone (PREG, Sigma-Aldrich) and lithium were administered subcutaneously (s.c.) at 2 ml per kg body weight once per day from gestational day (GD) 15 to post-natal day (PND) 23. PREG and lithium (Sigma-Aldrich) were dissolved in 20 % β-Cyclodextrin and sterile saline (0.9 % NaCl), respectively.

#### 2.2.2 Treatments

Primiparous female Sprague Dawley rats (Envigo) were used as mothers and single housed during pregnancy. Sprague Dawley dams expressing Cre recombinase under the control of the *TH* promoter (*TH*::Cre) were used for the DREADD experiments. THC or vehicle was administered (2 mg per kg, 2 ml per kg, s.c. once per day) from GD5 to GD20. This dose of THC was chosen because it does not produce behavioral responses or cannabinoid tolerance after repeated administration [45]. Moreover, this dose does not have any substantial impact on maternal and non-maternal behavior, as well as offspring bodyweight [36]. Additionally, this dose is equivalent to THC content in mild joint (5%) [46]. Offspring were weaned at PND 21 and were housed in a climate-controlled animal room (21 ± 1 °C; 60 % humidity) under a normal 12 h light-dark cycle (lights on at 7:00 a.m) with *ad libitum* access to water and food until the experimental day (PND15-28). To control for litters effects, we did not use more than two offsprings from each litter for the same experiment.

### 2.3 Surgical procedures

For chemogenetic manipulation, *TH*::Cre-positive offspring were stereotaxically bilaterally injected under isoflurane (4-5 % induction, 1-2 % maintenance) with a Cre-dependent adeno-associated virus expressing an inhibitory or excitatory DREADD construct, AAV5-hSyn-DIO-hM4D(Gi)-mCherry and AAV5-hSyn-DIO-hM3D(Gq)-mCherry, respectively, or control virus (AAV5-hSyn-DIO-mCherry) to target dopaminergic neurons in the VTA at PND7 (−4.2 mm posterior to bregma, ±0.55 mm lateral to bregma, −5.25 mm ventral from the cortical surface) with a Hamilton syringe. Viruses were injected at a volume of 0.5 μl per side at a rate of 0.1 μl/min. Injection needles were left in place for 5 min after the injection to ensure adequate viral delivery. For all the experiments the virus was incubated for at least 21 days, when expression was identifiable by the reporter protein expression.

### 2.4 Behavioral experiments

#### 2.4.1 Forced swim test and acute restraint stress

The forced swim test (FST) was conducted as previously described [44]. Briefly, rats were placed for 10 min in a transparent cylinder (40 cm high × 20 cm in diameter) filled with cold water to a depth of 30 cm, ensuring that the animals were unable to stabilize themselves by touching the bottom of the cylinder with their tails. After the FST, the rats were immediately dried out and then transferred to the startle cages for pre-pulse inhibition (PPI) testing.

Acute restraint stress (RS) was produced by placing the rat for 20 min in a well-ventilated, plastic tube (3.5 cm long × 6 cm in diameter) with an adjustable end to accommodate the size of each animal, as previously described [44]. As with the FST, after the RS, the animals were immediately subjected to the PPI procedure.

#### 2.4.2 Pre-pulse Inhibition test

Pre-pulse Inhibition (PPI) was tested following the protocol previously described [36]. The apparatus used to detect startle reflex and PPI parameters (Med Associates) consisted of four standard cages placed in sound-attenuated chambers with fan ventilation. Each cage consisted of a Plexiglas cylinder of 5 cm diameter, mounted on a piezoelectric accelerometric platform connected to an analog-digital converter. Two separate speakers conveyed background noise and acoustic bursts; each one properly placed so as to produce a variation of sound within 1 dB across the startle cage. On the testing day, each rat was placed in the cage for a 5-min acclimatization period consisting of 70-dB white noise background, which continued for the remainder of the session. Each session consisted of three consecutive sequences of trials (blocks). During the first and third blocks, rats were presented with only five pulse-alone trials of 115-dB. The second block consisted of a pseudorandom sequence of fifty trials, including twelve pulse-alone trials; thirty trials with a pulse preceded by 74-, 78-, or 82-dB prepulses (10 for each level of prepulse loudness); and eight no-stimulus trials, where only the background noise was delivered. Pulse and prepulse durations were set at 40 and 20 ms, respectively. Inter-trial intervals were selected randomly between 10 and 15 s, while the inter-stimulus intervals were set at 100 ms. The startle response was based on the first positive wave that meets the minimum wave criteria and determined as the mean startle amplitude of the pulse-alone trials relative to the second block. The % PPI was calculated only on the values relative to the second block using the following formula: [(mean startle amplitude for pulse alone trials - mean startle amplitude for prepulse + pulse trials)/mean startle amplitude for pulse alone trials] × 100. Depending on the experimental procedures, the PPI test was conducted either immediately after stress or 30 min after systemic administration of CNO to ensure the engagement of Gq/Gi-DREADDs in the VTA.

### 2.5 Immunohistochemistry and cell counting

#### 2.5.1 Tissue preparation

Following the behavioral experiments, rats at PND 26-34 were deeply anaesthetized with isoflurane and transcardially perfused with saline, followed by 4% paraformaldehyde in 0.1 M phosphate buffer (PB; pH=7.4). Afterwards, brains were removed, postfixed 2 h in the same solution at 4°C, then rinsed three times in PB saline 1× (PBS) and preserved in the same solution at 4⁰C. The next day, brains were coronally cut on a vibratome (VT1000S, Leica Biosystems) to yield sections (thickness, 40 μm) suited for immunohistochemistry (IHC) processing. For each rat, three coronal sections representative of the VTA were collected based on stereotaxic coordinates ranging from −4.80 mm to −5.80 mm relative to bregma. These coordinates were referenced from the rat brain atlas by Paxinos and Watson [47].

#### 2.5.2 Reaction protocol, image acquisition, and cell counting

Free-floating sections were rinsed in 0.1 M PB, blocked in a solution containing 10% normal goat serum (NGS, Vector, UK) and 0.5% Triton X-100 in 0.1 M PB at room temperature (2 h). Thereafter, sections were incubated at 4°C (48h) with the rabbit polyclonal primary antibody anti-TH (1:1000, Merck, Germany, #AB152), rinsed three times in 0.1 M PB, and then incubated with the secondary antibody, Atto® 488-labeled goat anti-rabbit IgG (1:400, Merck, Germany, #18772) in 0.1 M PB at room temperature (3 h). Afterward, sections were incubated for 10 minutes in 4’,6-diamidino-2-phenylindole (DAPI; 1:10,000, Merck, Italy, D9542), to allow visualization of cell nuclei, rinsed in PB 0.1 M, and mounted onto super-frost glass slides using Mowiol® mounting medium.

Images of single wavelength (14-bit depth) were obtained with a ZEISS Axio Scan Z1 slide scanner (Zeiss, Germany). Brain sections were captured at 20× magnification (Objective: Plan-Apochromat 20×/0.8 M27) to acquire the whole VTA from both hemispheres. The ImageJ software (National Institutes of Health, USA) was used to quantify the number of TH+, mCherry+, and TH+-mCherry+ cells located in the VTA. Specifically, images were first background-adjusted, then cells were manually counted within the VTA by using the multi-point tool. Analyses were performed blind with respect to the treatment received by each animal. No significant differences in the relative proportion of TH+, mCherry+, and TH+-mCherry+ cells were found among the three sections, therefore values from different antero-posterior levels were averaged. For each experimental condition, the final percentages were calculated as an average of the values from each rat within the same experimental group.

### 2.6 Corticosterone and Adrenocorticotropic hormone Analysis

Rats were sacrificed by decapitation and blood was immediately collected from the trunk into K3-EDTA tubes, then centrifuged at 900×g for 15 min at 4°C; the resulting plasma was collected and frozen at −20°C until assayed. An enzyme-linked immunosorbent assay (ELISA) was used to quantify plasma levels of Corticosterone (CORT; #RE52211 IBL Corticosterone Enzyme Immunoassay Kit, TECAN Europe), and Adrenocorticotropic hormone (ACTH; #EK-001-21, Phoenix Pharmaceuticals Inc., Burlingame, CA, USA), as previously described [48]. ELISA assays were performed according to the manufacturer’s instructions using a 96-well plate pre-coated with polyclonal antibodies against an antigenic site on the CORT or ACTH molecules, respectively. The kits also provided a seven-point standard curve ready to use (0 to 83.2 ng/ml, CORT) or a peptide standard for a six-point serial dilutions (0 to 25 ng/ml, ACTH), as well as quality controls. Each sample was run in duplicate. CORT and ACTH plasma levels are expressed in ng/ml.

### 2.7 Quantitative Real-Time PCR Analysis

To characterize the gene expression levels of MRs and GRs, animals were sacrificed by decapitation, the brain was rapidly removed, and bilateral VTA tissue punches were manually dissected and stored at −80°C until use. VTA samples were homogenized in a Douncer containing 1 mL of QIAzol Lysis Reagent and then mixed with 200 uL of chloroform. After centrifugation, the supernatant (aqueous phase) was transferred to a new tube, and the RNeasy Lipid Tissue Mini Kit (QIAGEN) was used according to the manufacturer’s instructions. Once the total RNA was isolated, the quality and concentration were assessed using a NanoDrop 1000 Spectrophotometer (Thermo Fisher Scientific) and immediately stored at −80°C until further use. Total RNA from each VTA sample was transcribed into complementary DNA (cDNA) using the High-Capacity cDNA Reverse Transcription Kit (Applied Biosystems, CA, United States) in a 20-µl reaction volume and stored at −20°C until use. Reverse transcriptase reactions were conducted at 25°C for 10 min, 2 h at 37°C, and 5 min at 85°C. The final cDNA concentration was normalized across samples to 30 ng/μL with autoclaved Milli-Q water.

The MRs and GRs were assessed by qRT-PCR analyzing the gene expression levels of the nuclear receptor subfamily 3, group C, member 2 (*Nr3c2*) and the nuclear receptor subfamily 3, group C, member 1 (*Nr3c1*), respectively. Primers were designed and verified using the primer-BLAST design tool (Table 1). All samples were tested in triplicate and β-actin (*Actb* gene) was used as an endogenous housekeeping gene to normalize the transcriptional levels of all target genes analyzed. qRT-PCR was performed in an Optical 384-well plate with a QuantStudio™ 12K Sequence Detection System (Applied Biosystems, CA, United States) consisting of 2 activation steps (50°C for 2 min, then 95°C for 10 min) followed by 45 cycles of melting (95°C for 15 s) and annealing (60 °C for 1 min). The PCR reaction contained PowerUp SYBR Green Master Mix (Applied Biosystems, CA, United States). The comparative cycle threshold (ΔΔCt) method was used to establish the gene expression relative quantification (RQ), and the results were reported as fold change compared with the CTRL group for each sex.

**Table 1.**
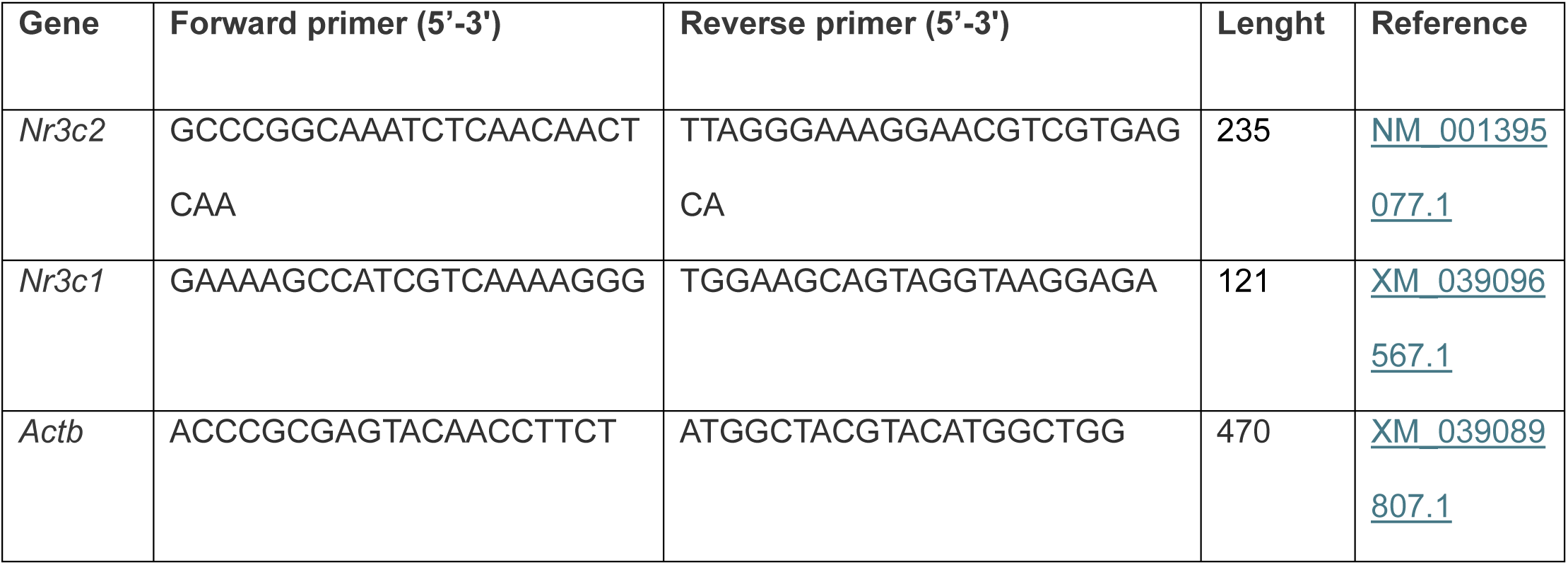
Primers for MRs and GRs characterization.

### 2.8 Statistical analysis

Rats were randomly assigned to each group. Statistical analysis was performed with GraphPad Prism 8 (San Diego CA, USA). Data were analyzed using 2- or 3-way ANOVA when appropriated, followed by Sidak’s or Tukey’s test. Significance threshold was set at p < 0.05. Data are presented as means ± SEM.

## 3 Results

### 3.1 Acute stress does not impair gating functions in PCE female progeny

Stress is one of the major determinants in the onset of dopamine-related psychopathologies [7, 8] and sex differences in stress-coping strategies have been already reported in experimental animals [7]. We previously found that PCE interferes with behavioral adaptations to an acute inescapable stress, such as the forced swim test (FST), in male [44] but not in female offspring [37]. FST also impairs subsequent PPI performance in PCE males [44]. To test whether PCE female rats were also protected by the effects of this acute inescapable stressor (i.e., FST) on PPI, PCE rats were subjected to FST before the PPI (Fig. 1A). FST induced sex-specific deficits in PPI, being CTRL females less performant than males (Fig. 1B; 2way ANOVA, F_1,43_= 5,21; p=0.0035). Of note, FST exerted sex-dependent PPI effects as a function of PCE (Fig. 1B; sex x PCE: 2way ANOVA, F_1,43_= 14,35; p=0.0005), supporting that PCE acts as a “first hit” [49] and endows the male offspring with an aberrant salient attribution [44], whereas the females display a normal behavioral performance [36, 37, 44]. To assess whether this reaction depends on a sex-specific engagement of the HPA axis in the PCE progeny, we measured baseline and stress-induced rise of endogenous plasma corticosterone (CORT) and adrenocorticotropic hormone (ACTH) levels. First, in a cohort of unstressed PCE rats, we verified whether PCE alters baseline CORT and ACTH levels as a function of sex (Fig. 1C-D). No differences were found among the groups (CORT: stress x PCE x sex: 3way ANOVA, F_1,60_=0,12; p=0.7303; ACTH: stress x PCE x sex: 3way ANOVA, F_1,60_=2,01; p=0.1613; Fig. 1C-D). Then, in line with previous findings [50], FST raised plasma CORT and ACTH concentrations to a larger extent in female rats irrespective of PCE (CORT: 2way ANOVA, F_1,32_=15,28; p=0.0005; ACTH: 2way ANOVA, F_1,32_=7,22; p=0.0113, Fig.1C-D). This suggests that plasma CORT and ACTH levels neither drive sex-specific coping strategies to FST [37, 44] nor stress-induced deficits in PPI progeny (Fig.1B and [44]).

**Figure 1.**
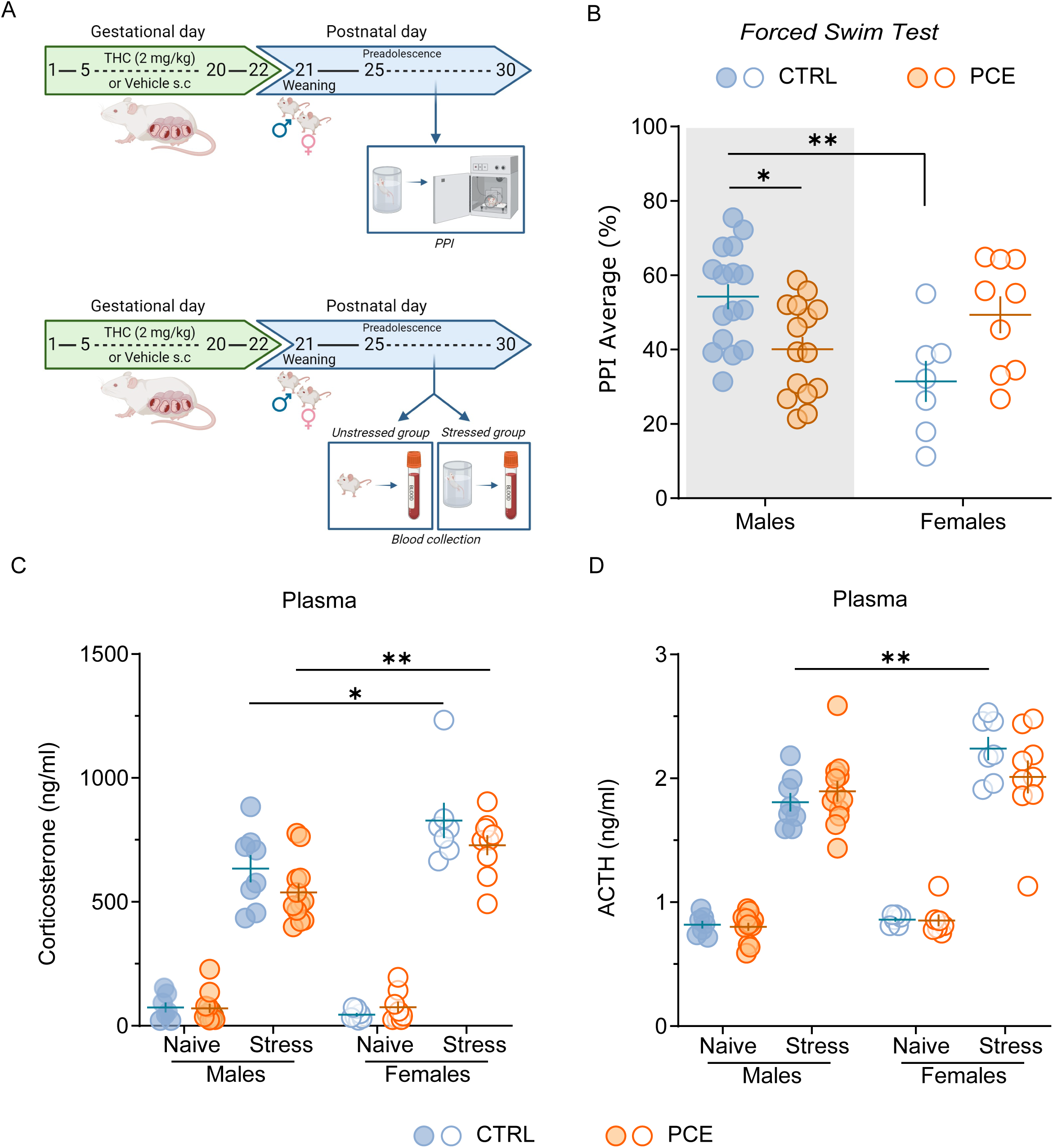
Sex-specific effects of acute stress on sensorimotor gating function. **A)** Timeline of experimental procedures for pre-pulse inhibition (PPI) experiments (top) and blood collection (bottom). **B)** Forced swim test (FST) differentially impacts on sensorimotor gating functions in male and female rats (*p = 0.036 CTRL males vs. PCE males; **p = 0.003 CTRL males vs. CTRL females; Sidak’s test; n_rats_ = 7-9 CTRL/PCE females, 15-16 CTRL/ PCE males). **C)** Corticosterone and **D)** adrenocorticotropic hormone (ACTH) concentrations were measured from plasma samples collected from unstressed rats and from stressed rats after 30 min from the beginning of the FST: *CORT*: *p = 0.01 stressed CTRL males vs. stressed CTRL females; **p = 0.003 stressed PCE males vs. stressed PCE females. *ACTH*: **p = 0.006 stressed CTRL males vs. stressed CTRL females; Sidak’s test. Data are expressed as ng/ml of plasma and obtained from 7-8 unstressed rats, 8-12 stressed. Unless otherwise indicated, data are represented with scatter plots with mean and ± SEM. Figure created with BioRender.com.

### 3.2 Sex-specific reduced feedback sensitivity to acute stress

Under stress conditions, CORT acts by activating high-affinity mineralocorticoid receptors (MRs) and low-affinity glucocorticoid receptors (GRs). MR/GR imbalance is suggested to compromise HPA axis responses to stress [7, 51]. Of note, CORT also acts locally in the VTA to modulate dopamine signaling [16, 52, 53]. To assess whether PCE induced an imbalance between MR- and GR-mediated actions, we determined mRNA levels of MR (*Nr3c2*) and GR (*Nr3c1*) in preadolescent unstressed and stressed rat VTA using quantitative real-time PCR (Fig 2A). Key differences were found with a specific unbalance between mRNA levels of *Nr3c2* (2way ANOVA, F_1,33_=7,20; p=0.0113; Fig.2B) and *Nr3c1* (2way ANOVA, F_1,44_=8,55; p=0.0054; Fig.2C) as a function of PCE only in unstressed male rats. FST further decreased mRNA levels of *Nr3c1* in PCE males (2way ANOVA, F_1,44_=4,12; p=0.04; Fig.2C), whereas it increased mRNA levels of *Nr3c2* in PCE females (*Nr3c2:* 2way ANOVA, F_1,25_=4,57; p=0.0425*; Nr3c1*: 2way ANOVA, F_1,29_=2,12; p=0.1558; Fig.2D-E), without modifying mRNA levels of *Nr3c2* (2way ANOVA, F_1,33_=0,01; p=0.89; Fig.2B) in PCE males and *Nr3c1* (2way ANOVA, F_1,44_=0,01; p=0.91; Fig. 2E) in females. Since MRs operate as an on-switch to select an appropriate coping response, whereas GRs serve as the off-switch to terminate the duration of the HPA-mediated reaction, collectively, this data suggests that male-specific MR/GR imbalances might accompany an inadequate HPA axis reaction to FST [44] and might explain the subsequent impaired PPI (Fig. 1B and [44]).

**Figure 2.**
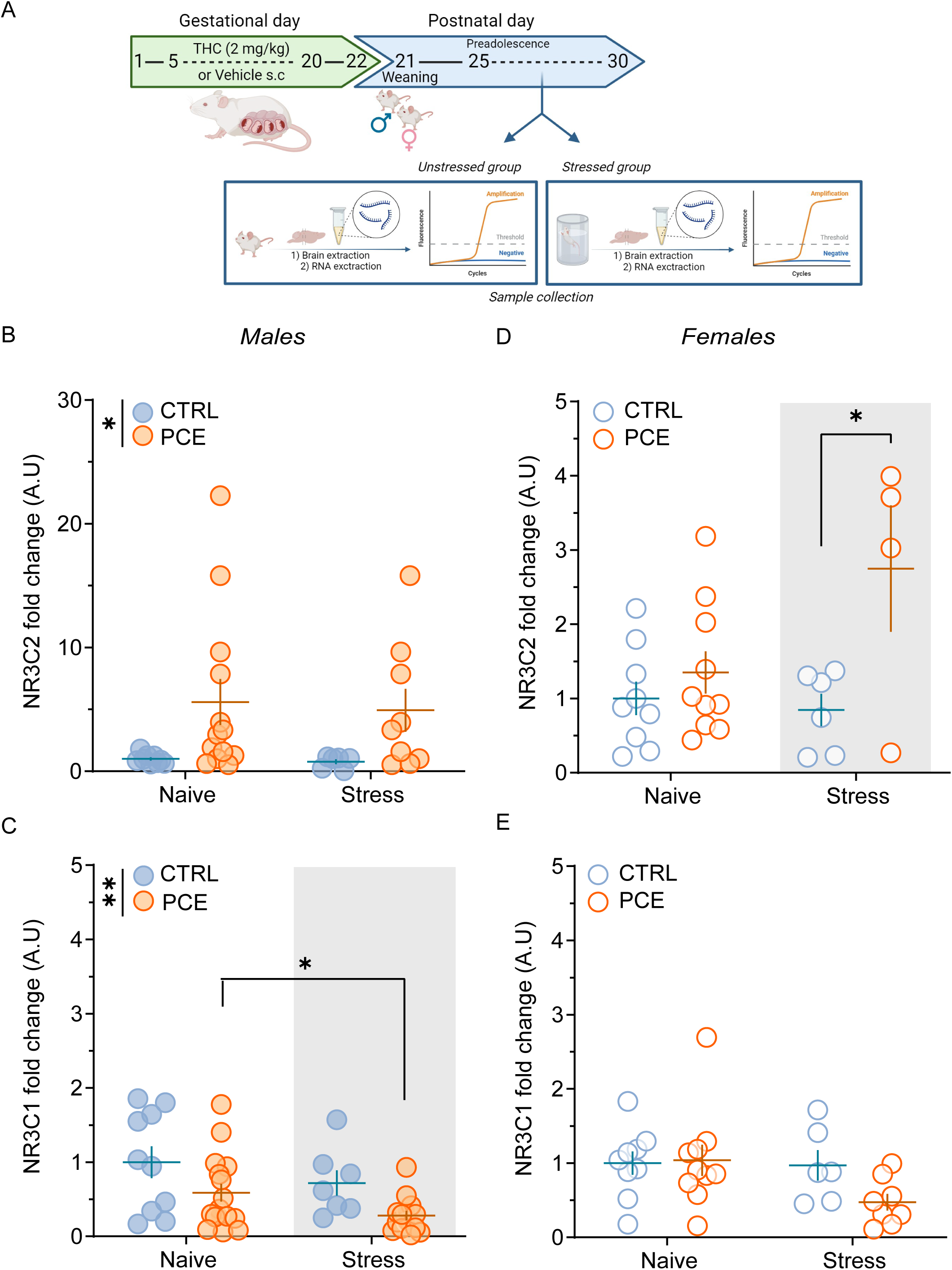
Feedback sensitivity to acute stress is reduced in PCE males. **A)** Timeline of experimental procedures. **B)** Fold change ratio of mRNA expression levels of Nr3c2 in VTA of males at preadolescence from unstressed rats and from stressed rats after the exposure to FST (*p = 0,0113 main effect of PCE; n_rats_ = 6-9 CTRL males, 9-13 PCE males). **C)** Graphs show the mRNA expression levels of Nr3c1 in VTA as a fold change ratio in male rats compared with their corresponding control group, either unstressed or stressed rats (**p = 0,0054 main effect of PCE; *p = 0,04 main effect of stress; n_rats_ = 7-10 CTRL males, 14-17 PCE males). **D)** Fold change ratio of mRNA expression levels of Nr3c2 in VTA of unstressed and stressed females at preadolescence (* p < 0.05 stressed PCE vs. stressed CTRL; Sidak’s test; n_rats_ = 6-9 CTRL females, 4-10 PCE females). **E)** Graphs show the mRNA expression levels of Nr3c1 in VTA as a fold change ratio in stressed and unstressed female rats compared with their corresponding control group (n_rats_ =6-9 CTRL females, 8-10 PCE females). Unless otherwise indicated, data are represented with scatter plots with mean and ± SEM. Figure created with BioRender.com.

### 3.3 Maladaptive responses to acute stress are mediated by dopamine

Dopamine role in the regulation of PPI has long been recognized [54–60] within an extremely complex circuit [41, 61]. However, the causative role of mesolimbic hyperdopaminergia underlying PPI deficits has never been tested by spatially isolating this single neural component. To test the hypothesis that stress-induced disruption of PPI in PCE males requires enhanced VTA dopamine signaling, we inhibited this cell population before FST. We induced the expression of Gi-coupled hM4D receptor in VTA dopamine neurons by injecting a Cre-dependent adeno-associated viral (AAV) vector, AAV5-hSyn-DIO-hM4D(Gi)-mCherry, into the VTA of male TH::Cre transgenic rats (Fig. 3A) in which Cre mimics the expression of the rate-limiting enzyme for dopamine synthesis, i.e., tyrosine hydroxylase (TH) [62]. Control rats received the AAV5-hSyn-DIO-mCherry vector, which carried the gene for mCherry alone. Within the VTA, 64±9.9 % of mCherry labelled cells were also TH-positive, whereas only 35.9±9.9 % of transduced cells were TH-negative (Fig. 3B), confirming the cell specificity of the DREADD (designer receptors exclusively activated by designer drugs) approach. Male rats were administered with clozapine N-oxide (CNO, 3 mg/kg i.p.) to inhibit VTA dopamine cells [63] before FST (Fig. 3C-D). Decreasing VTA dopamine neuron activity did not affect the startle amplitude (Fig. 3C; 2way ANOVA Gi-DREADD: F_1,78_ = 0,34; p=0.56), while it prevented FST-induced deficits of PPI in PCE male offspring (Fig. 3D; 2way ANOVA Gi-DREADD: F_1,35_= 0,35; p=0.56; 2way ANOVA Gi-DREADD x PCE: F_1,24_ = 11,7; p= 0.002 and [44]). In addition, inhibiting VTA dopamine signaling via chemogenetics prevented detrimental effects on PPI of another unavoidable stress condition, i.e. acute restraint stress (RS) in PCE male rats (Fig. 3F; 2way ANOVA Gi-DREADD: F_1,21_= 0,035; p=0.85) (2way ANOVA PCE: F_1,34_ = 6,17; p= 0.018; Fig. 3F) without affecting the startle amplitude (2way ANOVA Gi-DREADD: F_1,36_ = 4,427e-006, p=0.99; Fig. 3E). CNO is converted in clozapine and, therefore, may have off-target effects at other endogenous receptors rather than at Gi-DREADDs [64]. However, CNO administration in TH::Cre rats expressing a control fluorophore had no effect on both startle amplitude and PPI performance (Fig. 3C-F), thus indicating that the protective effects on PPI are DREADD-mediated and require normalizing VTA dopamine neuron activity.

**Figure 3.**
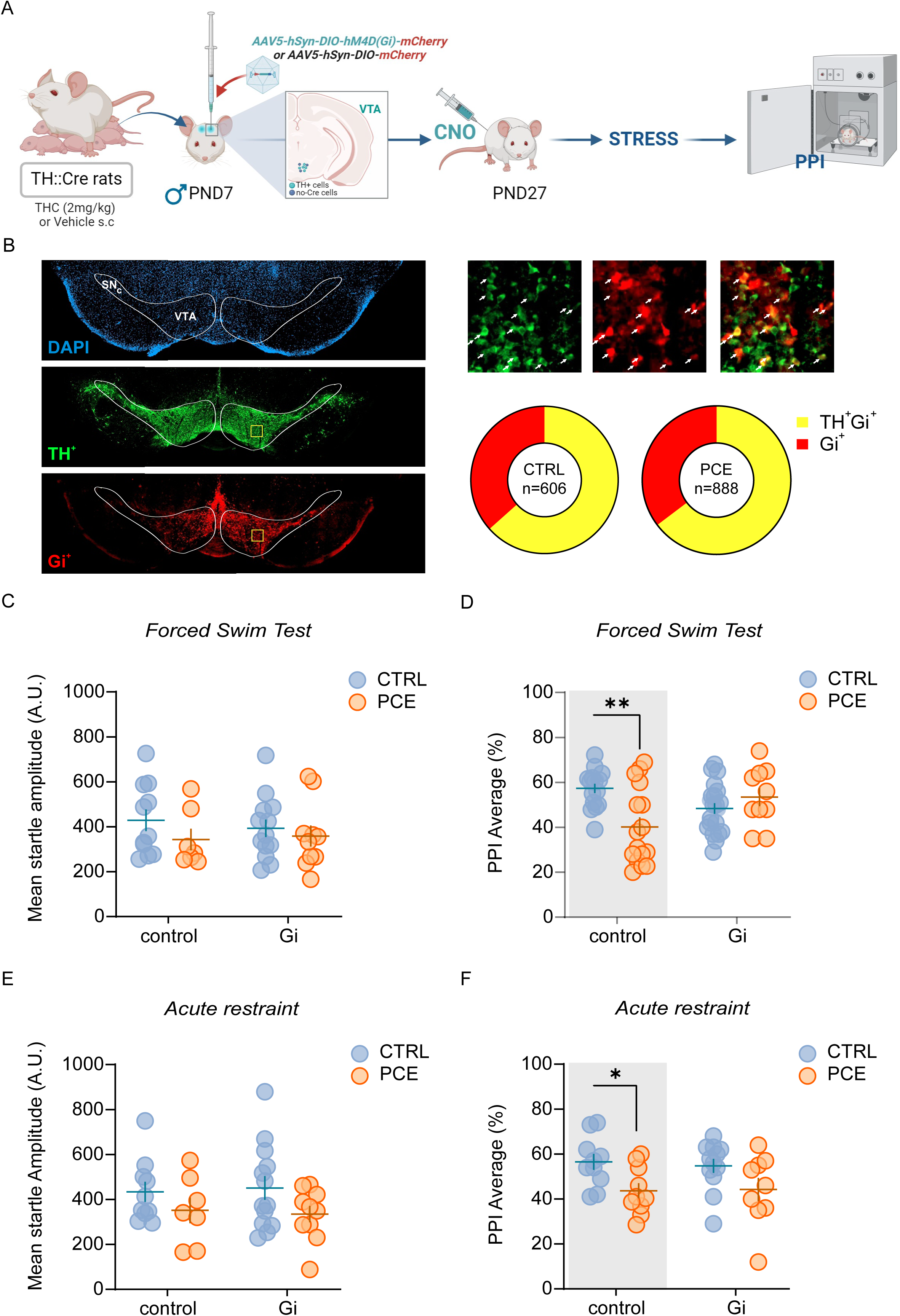
Maladaptive responses to acute stress are mediated by VTA dopaminergic activity. **A)** Schematic timeline of PCE treatment and behavioral experiments. **B)** Representative fluorescent images of Gi-mCherry and TH positive cells in coronal brain sections containing the VTA, counterstained for DAPI (left panel). Yellow square indicates one example ROI used for cell count (left panel). High magnification of yellow ROI for labelling quantification in VTA (top right): arrows indicate cells double positive for TH (green) and Gi-mCherry (red). Pie charts for cell quantification in CTRL and PCE animals (bottom right). **C)** Forced swim test (FST) does not affect the startle amplitude. Startle amplitude values are represented as arbitrary units (AU). (n_rats_ = 7 PCE-control; 10 PCE-Gi; 11 CTRL-control; 13 CTRL-Gi). **D)** Gi-DREADD activation prevents FST-induced deficits of sensorimotor gating functions in PCE males (**p=0.001 CTRL-control vs PCE-control; Sidak’s test; n_rats_ = 10 PCE-Gi; 16 CTRL-control and PCE-control; 21 CTRL-Gi). **E)** Effect of acute restraint stress (RS) in the startle amplitude after Gi-DREADD activation in PCE male offspring (n_rats_ = 10 CTRL-control and PCE-Gi; 13 CTRL-Gi; 7 PCE-control). **F)** Gi-DREADD activation prevents PPI deficits exhibited by PCE males (*p=0.04 CTRL-control vs PCE-control; Sidak’s test; n_rats_ = 9 PCE-Gi; 10 CTRL-control and PCE-control; 12 CTRL-Gi). Unless otherwise indicated, data are represented with scatter plots with mean and ± SEM. Figure created with BioRender.com.

To examine the pathological role of a hyperdopaminergic phenotype in PPI regulation by acute stress, we chemogenetically excited VTA DA cells [63] before FST in female rats, as their PPI performance is not impacted by this acute unescapable stressor as a function of PCE (Fig. 1B). We induced the expression of Gq-coupled hM3D receptor in VTA dopamine neurons by injecting a Cre-dependent AAV vector, AAV5-hSyn-DIO-hM3D(Gq)-mCherry, into the VTA of female TH::Cre transgenic rats (Fig. 4A). Control rats received the AAV5-hSyn-DIO-mCherry vector, which carried the gene for mCherry alone. Within the VTA, 73.9±10.1 % of mCherry labelled cells were also TH-positive, whereas only 26±10.2 % of transduced cells were TH-negative (Fig. 4B). Female rats were administered with CNO to stimulate VTA dopamine cells before FST and PPI (Fig. 4C). Importantly, a wide range of CNO doses, including 3 mg/kg, does not affect locomotor activity of this TH::Cre transgenic rat [65] but see Boekhoudt, Omrani, Luijendijk, Wolterink-Donselaar, Wijbrans, van der Plasse and Adan [66]. Unlike Gi-DREADD activation in males, in vivo excitation of VTA dopamine neuron activity by CNO markedly reduced startle amplitude in both CTRL and PCE females. This effect was observed after exposing animals to either FST or RS experimental procedures (Fig. 4C; FST: 2way ANOVA, Gq-DREADD, F_1,66_= 5.71, P<0.05; Fig. 4D; RS: 2way ANOVA, Gq-DREADD, F_1,61_= 65.89; P<0.0001). Importantly, no specific interactions among factors were detected (FST: 2way ANOVA, interaction Gq-DREADD x PCE, F_1,66_= 2.88, P=0.09, Fig. 4D; RS: 2way ANOVA, interaction Gq-DREADD x PCE, F_1,61_= 1.044; P=0.31, Fig. 4C), indicating that PCE did not significantly affect the decrement in startle amplitude produced by Gq-DREADD manipulation under both experimental stress conditions. Given that PPI data are extrapolated by startle values, and Gq-DREADD stimulation elicited a robust reduction in startle magnitude in both CTRL and PCE females, we computed %PPI to delta PPI to avoid potential artifacts in data interpretation caused by “floor effects” in startle magnitude. Delta PPI avoids this eventual flaw by calculating absolute differences between startle magnitudes on pulse-alone and prepulse+pulse trials [67–69]. Using this parameter, CNO significantly deteriorated PPI performance following FST and RS, irrespective of PCE (FST: 2way ANOVA, Gq-DREADD: F_1,66_= 4.23; P<0.05, Fig. 4E; RS: 2way ANOVA, Gq-DREADD: F_1,61_= 47.25; P=0.0001, Fig. 4F). Therefore, chemogenetic modulation of VTA dopamine neuronal activity is essential for the manifestation of stress-induced impairments of PPI in rats.

**Figure 4.**
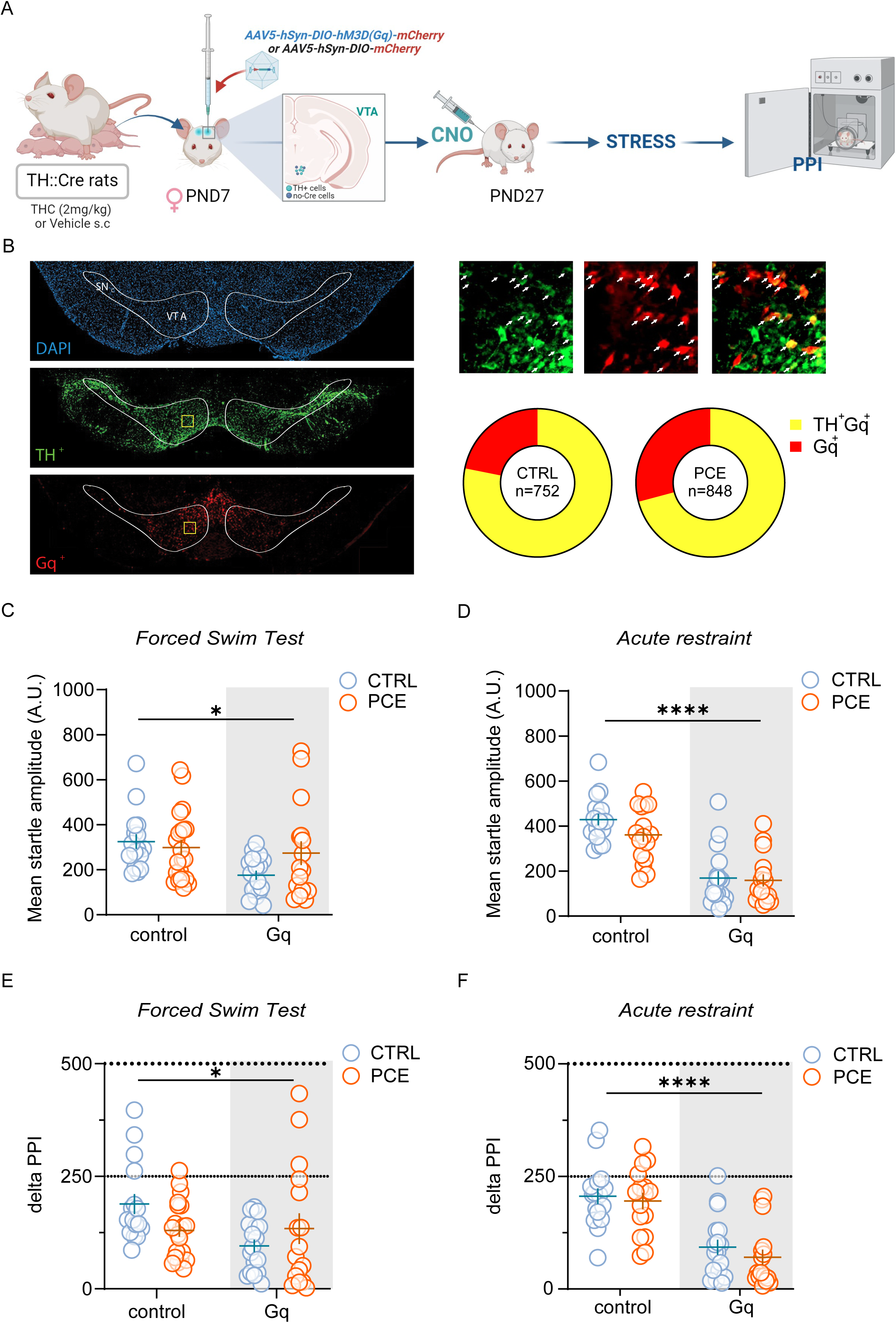
Chemogenetic activation of VTA DA neurons induces maladaptive responses to acute stress in female rats. **A)** Schematic timeline of PCE treatment and behavioral experiments. **B)** Representative fluorescent images of Gq-mCherry and TH positive cells in coronal brain sections containing the VTA, counterstained for DAPI (left panel). Yellow square indicates one example ROI used for cell count (left panel). High magnification of yellow ROI for labelling quantification in VTA (top right): arrows indicate cells double positive for TH (green) and Gq-mCherry (red). Pie charts for cell quantification in CTRL and PCE animals (bottom right). **C)** Gq-DREADD activation affects the startle amplitude in control female offspring after FST. Startle amplitude values are represented as arbitrary units (AU) (*p=0,019 main effect of Gq; n_rats_ = 16-17 per group). **D)** Effect of Gq-DREADD activation in the startle amplitude after the exposure to an acute restraint stress (RS) in female rats (**** p< 0.0001 main effect of Gq; n_rats_ = 16-17 per group). **E)** Gq-DREADD activation impairs sensorimotor gating functions following FST in female rats (*p=0,04 main effect of Gq; n_rats_ = 16-17 CTRL-control, CTRL-Gq, PCE-control; 21 PCE-Gq). **F)** Gq-DREADD activation induces PPI deficits in response to RS in female rats (**** p< 0.0001 main effect of Gq; n_rats_ = 16-17 per group). Unless otherwise indicated, data are represented with scatter plots with mean and ± SEM. Figure created with BioRender.com.

### 3.4 Pharmacological correction of stress-induced deficits of gating functions in PCE male progeny

We previously found that post-natal treatment with neurosteroid pregnenolone (PREG), FDA-approved for clinical trials, prevents PCE-induced hyperdopaminergic phenotypes and confers resilience against acute effects of THC in PCE male progeny [36]. To test whether PREG precludes stress-induced effects on PPI in PCE male progeny, we administered it (6 mg/kg s.c. once daily for 9 days, from post-natal day -PND-15 to 23) to CTRL and PCE male rats. Two days following the last administration, when PREG was cleared from the brain, we subjected the animals to FST and PPI (Fig. 5A). In agreement with our previous findings pointing to restorative actions of PREG towards dopamine system function [36], we did not observe stress-induced disruption of sensorimotor gating functions in PCE male offspring treated with PREG (2way ANOVA PCE x PREG: F_1,47_ = 8.803; P=0.0047, Fig. 5B). PREG treatment had no effect on PPI performance in CTRL male rats (CTRL-VEH vs. CTRL-PREG, P=0.68). PREG can also act by inhibiting the GSK3β [70], which is a key element in the pathogenesis of schizophrenia [71], and also a target of lithium [72, 73]. We, therefore, tested the hypothesis that PREG molecular target was GSK3β and that a postnatal treatment with lithium could mimic PREG effects on stress-induced deficits of PPI in PCE males. Lithium was administered to CTRL and PCE male rats (50 mg/kg s.c., once daily for 9 days, from PND 15 to 23; Fig. 5C), and 48 hours following the last administration, we subjected the animals to FST and PPI. Notably, lithium rescued PCE males from FST-induced deficits without affecting CTRL PPI performance (Fig. 5D; 2way ANOVA PCE x Lithium: F_1,28_ = 5,100; p=0.03).

**Figure 5.**
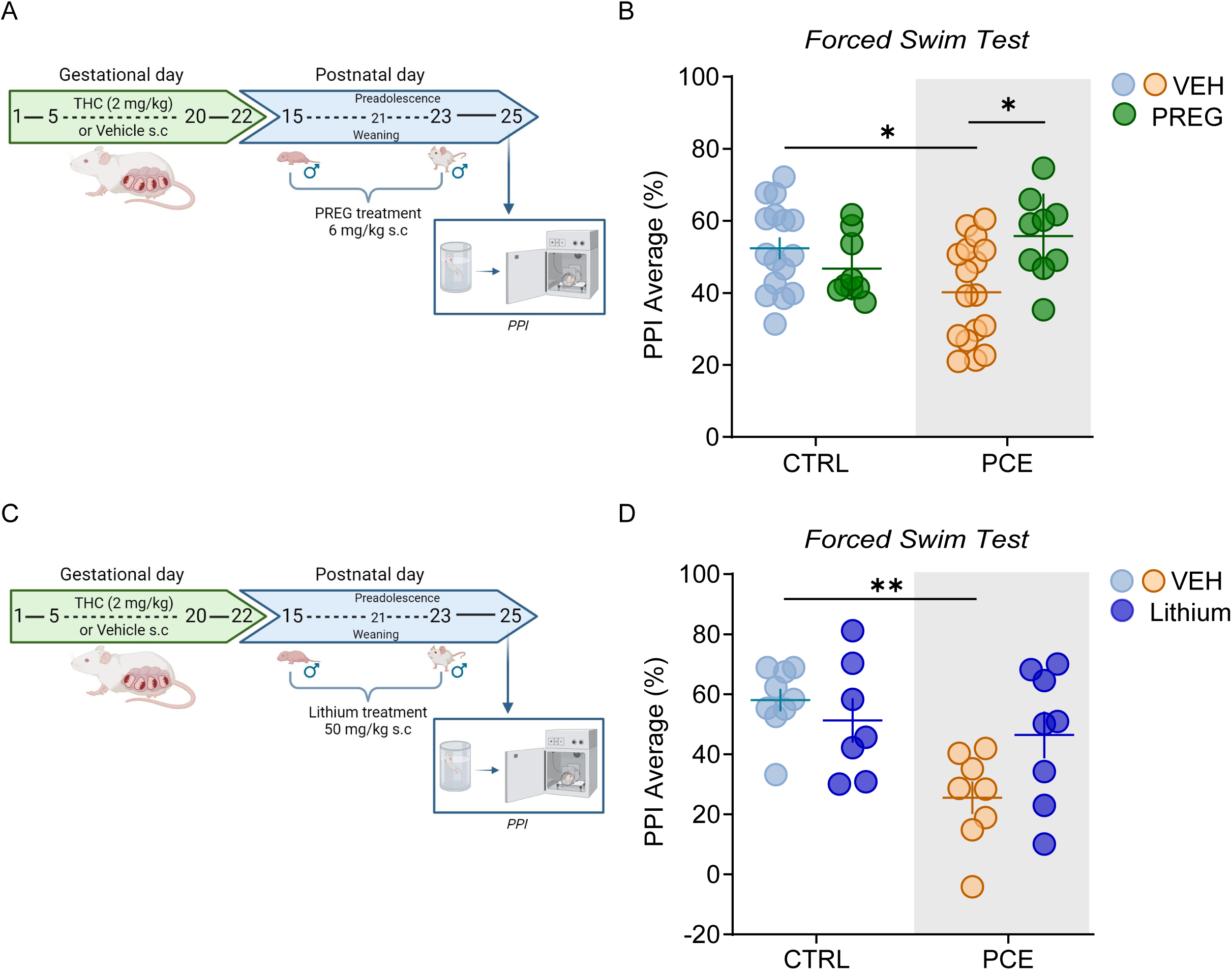
Pharmacological manipulations with pregnenolone or lithium prevent stress-induced deficits of gating functions in PCE male progeny. **A)** Schematic timeline of PCE treatment and subchronic treatment with pregnenolone (PREG). **B)** PREG prevents FST-induced deficits of PPI in male PCE offspring (*p=0.02 PCE-VEH vs. CTRL-VEH; *p=0.01 PCE-VEH vs. PCE-PREG; Tukey’s test; n_rats_ = 9 CTRL/PCE-PREG; 16-17 CTRL/PCE-VEH). **C)** Schematic timeline of PCE treatment and subchronic treatment with lithium. D) Lithium prevents FST-induced deficits of PPI in male PCE offspring (**p=0.003 PCE-VEH vs. CTRL-VEH; Tukey’s test; n_rats_ = 7-8 CTRL/PCE-Lithium; 8-9 CTRL/PCE-VEH). Unless otherwise indicated, data are represented with scatter plots with mean and ± SEM. Figure created with BioRender.com.

## 4. Discussion

In the present study, we demonstrate the essential role of dopamine in the detrimental effects of acute unescapable stressors on sensorimotor gating functions in preadolescent rats. DREADD-driven changes of VTA dopamine cell activity before the exposure to an acute inescapable stressor impact on subsequent PPI performance. This highlights the causal link between VTA dopamine signaling and the engagement of HPA axis for the individual ability to inhibit behavioral responses to incoming sensory information. As PCE promotes a male-specific dysregulation of dopamine and HPA axis responses within the VTA (present data and [36, 37]), such stress- and dopamine-induced impairment of sensorimotor gating functions only manifest in the male progeny. This is relevant because PCE children exhibit proneness to psychotic-like experiences and some psychopathological phenotypes where males face a higher risk [20, 23, 28, 74–76].

Our data support and extend previous findings suggesting that any manipulation leading to enhanced presynaptic dopamine release and/or dopamine receptor stimulation in the target region of the Nucleus Accumbens (NAc) disrupts PPI [55, 60, 77–81]. Of note, an association exists between hyper- and hypo-activation of mesolimbic and mesocortical dopamine pathways, respectively, and the inability of filter out relevant environmental stimuli to control adaptive motor behavior [56, 82]. Since a bidirectional chemogenetic modulation of dopamine neuronal activity in TH::Cre rats also modifies dopamine release in the NAc [63], it is plausible that CNO by engaging either Gi- or Gq-DREADD corrects or produces, respectively, a mesolimbic hyperdopaminergia, a signature of disrupted sensorimotor gating functions [38–41]. Our findings are in line with the evidence that hyperdopaminergic resting-states of VTA dopamine cells are necessary in the pathophysiology of psychosis and schizophrenia [83–86], and that chemogenetic approaches aimed at normalizing rather than silencing VTA dopamine neuronal activity prevent the expression of aberrant behaviors in animal models of schizophrenia [85, 86]. Accordingly, PCE in rodents produces male-specific hyper-dopaminergic neurophysiological phenotypes [36, 37, 87, 88], shifts the proportion of tonically firing dopamine cells [44] and the balance between excitation and inhibition [36, 88], which are collectively risk factors for susceptibility to diverse neuropsychiatric disorders. Since acute stress activates dopamine system function, which contributes to homeostatic neurobehavioral stress responses also by altering its sensitivity to subsequent stimuli [8, 12–14, 16, 52, 89, 90], PCE-induced sex-specific derangement of dopamine neuron resting-states may be critical for subsequent susceptibility to develop psychopathological phenotypes later in life [28, 91–96]. Of note, PCE produces enduring remarkable repercussions in the offspring that go beyond the brain-wide adaptations, including alterations in lipidomic and cytokine/chemokine profiles [88, 93, 94, 97–99], and extend from cardiac and metabolic dysfunction to changes in intestinal microbiota composition [98, 100]. Since gut microbiota couples immune responses to stress-sensitive brain circuits [101], these findings warrant further investigations to disentangle the mechanisms underpinning PCE sequelae on the progeny and raise concerns on whether these aberrant stress-dependent neurobehavioral reactions (present data and [37, 44]) relate to PCE impact on gut microbiota, lipid and/or glucose metabolism [88, 98], and/or immune system [97], and whether they do subside with development.

While the mechanisms underlying PCE sex-specific reduced feedback sensitivity to an acute stressor remain to be fully elucidated, our observations support the notion that an efficient on/off switch mediated by CORT via MR/GR activation is crucial for successful coping and adapting strategies to inescapable stressful situations [7, 102], thereby providing energy for resilience. Indeed, the observation that, in the VTA, PCE male rats only display enhanced and decreased gene expression levels of MRs and GRs, respectively, might explain the sex-specific failure to adopt coping/adaptive strategies during the FST [37, 44]. Accordingly, a less efficient termination of the HPA axis activity mediated by GRs, which operate as the off-switch, promotes energy expenditure and exaggerated behavioral responses [7, 51]. Conversely, our finding that acute FST only in PCE females increases gene expression levels of MRs, without affecting those of GRs, supports the notion that a gain-of-function of MRs combined with an efficient GR-dependent off-switch is associated with a “resilient” phenotype better equipped for acute stress responses of the HPA axis and for long-term adaptation [7, 51]. Accordingly, while acute stress increases plasma CORT and ACTH levels to a larger extent in females when compared to males (present data and [50]), and CORT is detrimental to PPI performance [103] (Fig. 1b), acute FST does not affect PPI in PCE female rats, which also show appropriate coping strategies during the FST [37]. Consequently, PCE-induced male-specific MR/GR imbalances within the VTA may take part to those metaplastic changes setting a resting state prone to maladaptive stress reaction and PPI deterioration, the latter requiring a hyperdopaminergic neurophysiological phenotype.

Finally, our study helps elucidating the mechanisms participating in sex-dependent PCE detrimental effects on PPI performance by establishing that these behavioral effects depend on abnormal GPCR signaling cascade engaged in VTA dopamine neurons, including GSK-3β, a molecular signature of diverse psychiatric conditions [71, 104–107]. Our results showing that PREG, which also inhibits the GSK-3β [108], corrects FST-induced deficits of PPI in PCE male offspring extend previous evidence that it can restore VTA dopamine circuit function and its responsiveness to an acute challenge of THC in PCE males [36]. Since PREG restorative effects are manifest in its absence (present data and [36]), cannot be ascribed to its well-known downstream metabolites (e.g., progesterone) [36], and are mimicked by lithium, our data suggests, among the others, GSK-3β as a possible molecular target candidate. Given that Gi-DREADD stimulation in VTA dopamine cells of PCE male rats rescues the aberrant phenotype, one might speculate the occurrence of a molecular convergence that involves hyperactivity of GSK-3β as an adaptive response to PCE. This is in line with the role of GSK3β in neurodevelopment [109], in the pathogenesis of diverse psychiatric disorders (mood disorders, schizophrenia, drug abuse)[106, 107, 110–112] and in response to medication [72, 113–115]. Thus, inhibiting its activity is considered of therapeutic interest.

As cannabis use among pregnant women has been on the rise [30, 31], our study deepens the knowledge on PCE detrimental effects on neurodevelopment [26, 75] and warrants further investigations on the extensive spectrum of neurobehavioral maladaptations induced by the different ingredients of cannabis and at different developmental stages to uncover age- and gender-specific therapeutic approaches.

## 5. Conclusions

Given the well-established interplay between stressful life events and the onset of psychopathologies, particularly during (pre)adolescence, our study helps delineating neuroendocrine mechanisms connecting, for the first time to our knowledge, stress and dopamine-dependent psychopathologies.

## 6. Acknowledgements

The authors thank S. Aramo, and the CeSASt personnel for their skillful assistance.

## 7. Author contributions

V.S., F.T., and L.C. performed behavioral experiments and analyzed the data. V.S., S.A., and R.L. performed the surgeries. P.P. and L.P-V. carried out the molecular experiments and analyzed the data. A.B-G., P.P. and R.F. analyzed molecular and behavioral data, and reviewed the manuscript. M.S. and R.L. carried out and analyzed immunohistochemical experiments. V.S. and R.L. prepared the figures. M.M. conceived, designed and supervised the project, provided funding and wrote the original draft.

## References

[1] GBD 2017 Disease and Injury Incidence and Prevalence Collaborators. Global, regional, and national incidence, prevalence, and years lived with disability for 354 diseases and injuries for 195 countries and territories, 1990–2017: a systematic analysis for the Global Burden of Disease Study 2017. The Lancet (2018).

[2] M. Solmi, J. Radua, M. Olivola, E. Croce, L. Soardo, G. Salazar de Pablo, J. Il Shin, J.B. Kirkbride, P. Jones, J.H. Kim, J.Y. Kim, A.F. Carvalho, M.V. Seeman, C.U. Correll, P. Fusar-Poli, Age at onset of mental disorders worldwide: large-scale meta-analysis of 192 epidemiological studies, Mol Psychiatry 27(1) (2022) 281–295.

[3] A. Caspi, R.M. Houts, A. Ambler, A. Danese, M.L. Elliott, A. Hariri, H. Harrington, S. Hogan, R. Poulton, S. Ramrakha, L.J.H. Rasmussen, A. Reuben, L. Richmond-Rakerd, K. Sugden, J. Wertz, B.S. Williams, T.E. Moffitt, Longitudinal Assessment of Mental Health Disorders and Comorbidities Across 4 Decades Among Participants in the Dunedin Birth Cohort Study, JAMA Netw Open 3(4) (2020) e203221.

[4] W.H. Organization, The World Health Report: 2001: Mental Health: New Understanding, New Hope. World Health Organization., (2001).

[5] I. Pereira-Figueiredo, E. Umeoka, Stress: Influences and Determinants of Psychopathology., Encyclopedia 4(2) (2024) 1026–1043.

[6] E. Nederhof, F.V. van Oort, E.M. Bouma, O.M. Laceulle, A.J. Oldehinkel, J. Ormel, Predicting mental disorders from hypothalamic-pituitary-adrenal axis functioning: a 3-year follow-up in the TRAILS study, Psychol Med 45(11) (2015) 2403–12.

[7] E.R. de Kloet, M. Joels, The cortisol switch between vulnerability and resilience, Mol Psychiatry 29(1) (2024) 20–34.

[8] M.A. Bloomfield, R.A. McCutcheon, M. Kempton, T.P. Freeman, O. Howes, The effects of psychosocial stress on dopaminergic function and the acute stress response, Elife 8 (2019).

[9] J.H. Baik, Stress and the dopaminergic reward system, Exp Mol Med 52(12) (2020) 1879–1890.

[10] H.C. Meyer, F.S. Lee, Translating Developmental Neuroscience to Understand Risk for Psychiatric Disorders, Am J Psychiatry 176(3) (2019) 179–185.

[11] M. Mandy, M. Nyirenda, Developmental Origins of Health and Disease: the relevance to developing nations, Int Health 10(2) (2018) 66–70.

[12] E.H. Douma, E.R. de Kloet, Stress-induced plasticity and functioning of ventral tegmental dopamine neurons, Neurosci Biobehav Rev 108 (2020) 48–77.

[13] E.N. Holly, K.A. Miczek, Ventral tegmental area dopamine revisited: effects of acute and repeated stress, Psychopharmacology (Berl) 233(2) (2016) 163–86.

[14] O. Valenti, D.J. Lodge, A.A. Grace, Aversive stimuli alter ventral tegmental area dopamine neuron activity via a common action in the ventral hippocampus, J Neurosci 31(11) (2011) 4280–9.

[15] J.L. Niehaus, M. Murali, J.A. Kauer, Drugs of abuse and stress impair LTP at inhibitory synapses in the ventral tegmental area, Eur J Neurosci 32(1) (2010) 108–17.

[16] D. Saal, Y. Dong, A. Bonci, R.C. Malenka, Drugs of abuse and stress trigger a common synaptic adaptation in dopamine neurons, Neuron 37(4) (2003) 577–82.

[17] J.C. Pruessner, F. Champagne, M.J. Meaney, A. Dagher, Dopamine release in response to a psychological stress in humans and its relationship to early life maternal care: a positron emission tomography study using [11C]raclopride, J Neurosci 24(11) (2004) 2825–31.

[18] M. Colizzi, A. Lasalvia, M. Ruggeri, Prevention and early intervention in youth mental health: is it time for a multidisciplinary and trans-diagnostic model for care?, Int J Ment Health Syst 14 (2020) 23.

[19] K. Bolhuis, M.E. Koopman-Verhoeff, L.M.E. Blanken, D. Cibrev, V.W.V. Jaddoe, F.C. Verhulst, M.H.J. Hillegers, S.A. Kushner, H. Tiemeier, Psychotic-like experiences in pre-adolescence: what precedes the antecedent symptoms of severe mental illness?, Acta Psychiatr Scand 138(1) (2018) 15–25.

[20] J.D. Fine, A.L. Moreau, N.R. Karcher, A. Agrawal, C.E. Rogers, D.M. Barch, R. Bogdan, Association of Prenatal Cannabis Exposure With Psychosis Proneness Among Children in the Adolescent Brain Cognitive Development (ABCD) Study, JAMA Psychiatry (2019).

[21] S.E. Paul, A.S. Hatoum, J.D. Fine, E.C. Johnson, I. Hansen, N.R. Karcher, A.L. Moreau, E. Bondy, Y. Qu, E.B. Carter, C.E. Rogers, A. Agrawal, D.M. Barch, R. Bogdan, Associations Between Prenatal Cannabis Exposure and Childhood Outcomes: Results From the ABCD Study, JAMA Psychiatry (2020).

[22] S. Singh, K.B. Filion, H.A. Abenhaim, M.J. Eisenberg, Prevalence and outcomes of prenatal recreational cannabis use in high-income countries: a scoping review, BJOG 127(1) (2020) 8–16.

[23] A.W. Tadesse, G. Ayano, B.A. Dachew, K. Betts, R. Alati, Exposure to maternal cannabis use disorder and risk of autism spectrum disorder in offspring: A data linkage cohort study, Psychiatry Res 337 (2024) 115971.

[24] M.M. Faraj, J. Evanski, C.G. Zundel, C. Peters, S. Brummelte, L. Lundahl, H.A. Marusak, Impact of prenatal cannabis exposure on functional connectivity of the salience network in children, J Neurosci Res 101(1) (2023) 162–171.

[25] N.D. Volkow, W.M. Compton, E.M. Wargo, The Risks of Marijuana Use During Pregnancy, Jama 317(2) (2017) 129–130.

[26] A.W. Tadesse, B.A. Dachew, G. Ayano, K. Betts, R. Alati, Prenatal cannabis use and the risk of attention deficit hyperactivity disorder and autism spectrum disorder in offspring: A systematic review and meta-analysis, J Psychiatr Res 171 (2024) 142–151.

[27] R.D. Eiden, S. Shisler, D.A. Granger, P. Schuetze, J. Colangelo, M.A. Huestis, Prenatal Tobacco and Cannabis Exposure: Associations with Cortisol Reactivity in Early School Age Children, Int J Behav Med 27(3) (2020) 343–356.

[28] G. Rompala, Y. Nomura, Y.L. Hurd, Maternal cannabis use is associated with suppression of immune gene networks in placenta and increased anxiety phenotypes in offspring, Proc Natl Acad Sci U S A 118(47) (2021).

[29] L.R. Stroud, G.D. Papandonatos, N.C. Jao, C. Vergara-Lopez, M.A. Huestis, A.L. Salisbury, Prenatal tobacco and marijuana co-use: Sex-specific influences on infant cortisol stress response, Neurotoxicol Teratol 79 (2020) 106882.

[30] S. Hayes, E. Delker, G. Bandoli, The prevalence of cannabis use reported among pregnant individuals in the United States is increasing, 2002-2020, J Perinatol 43(3) (2023) 387–389.

[31] Q.L. Brown, A.L. Sarvet, D. Shmulewitz, S.S. Martins, M.M. Wall, D.S. Hasin, Trends in Marijuana Use Among Pregnant and Nonpregnant Reproductive-Aged Women, 2002-2014, Jama 317(2) (2017) 207–209.

[32] N.D. Volkow, B. Han, W.M. Compton, E.F. McCance-Katz, Self-reported Medical and Nonmedical Cannabis Use Among Pregnant Women in the United States, Jama 322(2) (2019) 167–169.

[33] B. Dickson, C. Mansfield, M. Guiahi, A.A. Allshouse, L.M. Borgelt, J. Sheeder, R.M. Silver, T.D. Metz, Recommendations From Cannabis Dispensaries About First-Trimester Cannabis Use, Obstet Gynecol 131(6) (2018) 1031–1038.

[34] M. O’Connor, Medicinal Cannabis in Pregnancy - Panacea or Noxious Weed?, J Law Med 25(3) (2018) 634–646.

[35] M. Besse, K. Parikh, K. Mark, Reported Reasons for Cannabis Use Before and After Pregnancy Recognition, J Addict Med 17(5) (2023) 563–567.

[36] R. Frau, V. Miczán, F. Traccis, S. Aroni, C.I. Pongor, P. Saba, V. Serra, C. Sagheddu, S. Fanni, M. Congiu, P. Devoto, J.F. Cheer, I. Katona, M. Melis, Prenatal THC produces a hyperdopaminergic phenotype rescued by pregnenolone, Nat Neurosci 22(12) (2019) 11.

[37] F. Traccis, V. Serra, C. Sagheddu, M. Congiu, P. Saba, G. Giua, P. Devoto, R. Frau, J.F. Cheer, M. Melis, Prenatal THC Does Not Affect Female Mesolimbic Dopaminergic System in Preadolescent Rats, Int J Mol Sci 22(4) (2021).

[38] J.G. Howland, E.M. MacKenzie, T.T. Yim, P. Taepavarapruk, A.G. Phillips, Electrical stimulation of the hippocampus disrupts prepulse inhibition in rats: frequency- and site-dependent effects, Behav Brain Res 152(2) (2004) 187–97.

[39] J.E. Lisman, J.T. Coyle, R.W. Green, D.C. Javitt, F.M. Benes, S. Heckers, A.A. Grace, Circuit-based framework for understanding neurotransmitter and risk gene interactions in schizophrenia, Trends Neurosci 31(5) (2008) 234–42.

[40] D.J. Lodge, A.A. Grace, Aberrant hippocampal activity underlies the dopamine dysregulation in an animal model of schizophrenia, The Journal of neuroscience : the official journal of the Society for Neuroscience 27(42) (2007) 11424–30.

[41] N.R. Swerdlow, G.A. Light, Sensorimotor gating deficits in schizophrenia: Advancing our understanding of the phenotype, its neural circuitry and genetic substrates, Schizophr Res 198 (2018) 1–5.

[42] J.P. Vargas, E. Diaz, M. Portavella, J.C. Lopez, Animal Models of Maladaptive Traits: Disorders in Sensorimotor Gating and Attentional Quantifiable Responses as Possible Endophenotypes, Front Psychol 7 (2016) 206.

[43] M.A. Geyer, K. Krebs-Thomson, D.L. Braff, N.R. Swerdlow, Pharmacological studies of prepulse inhibition models of sensorimotor gating deficits in schizophrenia: a decade in review, Psychopharmacology (Berl) 156(2-3) (2001) 117–54.

[44] C. Sagheddu, F. Traccis, V. Serra, M. Congiu, R. Frau, J.F. Cheer, M. Melis, Mesolimbic dopamine dysregulation as a signature of information processing deficits imposed by prenatal THC exposure, Prog Neuropsychopharmacol Biol Psychiatry 105 (2021) 110128.

[45] J.L. Wiley, M. O’Connell M, M.E. Tokarz, M.J. Wright, Jr., Pharmacological effects of acute and repeated administration of Delta(9)-tetrahydrocannabinol in adolescent and adult rats, J Pharmacol Exp Ther 320(3) (2007) 1097–105.

[46] Z. Mehmedic, S. Chandra, D. Slade, H. Denham, S. Foster, A.S. Patel, S.A. Ross, I.A. Khan, M.A. ElSohly, Potency trends of Delta9-THC and other cannabinoids in confiscated cannabis preparations from 1993 to 2008, J Forensic Sci 55(5) (2010) 1209–17.

[47] W.C. Paxinos G, The rat brain in stereotaxic coordinates, 6th Edition ed., Academic Press, San Diego, CA, 2007.

[48] G. Boero, M.G. Pisu, F. Biggio, L. Muredda, G. Carta, S. Banni, E. Paci, P. Follesa, A. Concas, P. Porcu, M. Serra, Impaired glucocorticoid-mediated HPA axis negative feedback induced by juvenile social isolation in male rats, Neuropharmacology 133 (2018) 242–253.

[49] K.A. Richardson, A.K. Hester, G.L. McLemore, Prenatal cannabis exposure - The “first hit” to the endocannabinoid system, Neurotoxicol Teratol 58 (2016) 5–14.

[50] M.B. Solomon, M. Loftspring, A.D. de Kloet, S. Ghosal, R. Jankord, J.N. Flak, A.C. Wulsin, E.G. Krause, R. Zhang, T. Rice, J. McKlveen, B. Myers, J.G. Tasker, J.P. Herman, Neuroendocrine Function After Hypothalamic Depletion of Glucocorticoid Receptors in Male and Female Mice, Endocrinology 156(8) (2015) 2843–53.

[51] A.P. Harris, M.C. Holmes, E.R. de Kloet, K.E. Chapman, J.R. Seckl, Mineralocorticoid and glucocorticoid receptor balance in control of HPA axis and behaviour, Psychoneuroendocrinology 38(5) (2013) 648–58.

[52] S.S. Daftary, J. Panksepp, Y. Dong, D.B. Saal, Stress-induced, glucocorticoid-dependent strengthening of glutamatergic synaptic transmission in midbrain dopamine neurons, Neurosci Lett 452(3) (2009) 273–6.

[53] A.R. de Oliveira, A.E. Reimer, M.L. Brandao, Mineralocorticoid receptors in the ventral tegmental area regulate dopamine efflux in the basolateral amygdala during the expression of conditioned fear, Psychoneuroendocrinology 43 (2014) 114–25.

[54] S.B. Powell, X. Zhou, M.A. Geyer, Prepulse inhibition and genetic mouse models of schizophrenia, Behav Brain Res 204(2) (2009) 282–94.

[55] R.S. Mansbach, M.A. Geyer, D.L. Braff, Dopaminergic stimulation disrupts sensorimotor gating in the rat, Psychopharmacology (Berl) 94(4) (1988) 507–14.

[56] D.L. Braff, M.A. Geyer, Sensorimotor gating and schizophrenia. Human and animal model studies, Arch Gen Psychiatry 47(2) (1990) 181–8.

[57] J.M. Doherty, V.L. Masten, S.B. Powell, R.J. Ralph, D. Klamer, M.J. Low, M.A. Geyer, Contributions of dopamine D1, D2, and D3 receptor subtypes to the disruptive effects of cocaine on prepulse inhibition in mice, Neuropsychopharmacology 33(11) (2008) 2648–56.

[58] J. Zhang, C. Forkstam, J.A. Engel, L. Svensson, Role of dopamine in prepulse inhibition of acoustic startle, Psychopharmacology (Berl) 149(2) (2000) 181–8.

[59] C. Nasello, L.A. Poppi, J. Wu, T.F. Kowalski, J.K. Thackray, R. Wang, A. Persaud, M. Mahboob, S. Lin, R. Spaseska, C.K. Johnson, D. Gordon, F. Tissir, G.A. Heiman, J.A. Tischfield, M. Bocarsly, M.A. Tischfield, Human mutations in high-confidence Tourette disorder genes affect sensorimotor behavior, reward learning, and striatal dopamine in mice, Proc Natl Acad Sci U S A 121(19) (2024) e2307156121.

[60] C. Yang, X. Chen, J. Xu, W. Chen, SKF82958, a dopamine D1 receptor agonist, disrupts prepulse inhibition in the medial prefrontal cortex and nucleus accumbens in C57BL/6J mice, Behav Pharmacol 35(4) (2024) 193–200.

[61] B. Acevedo, E. Aron, S. Pospos, D. Jessen, The functional highly sensitive brain: a review of the brain circuits underlying sensory processing sensitivity and seemingly related disorders, Philos Trans R Soc Lond B Biol Sci 373(1744) (2018).

[62] A.J. Brown, D.A. Fisher, E. Kouranova, A. McCoy, K. Forbes, Y. Wu, R. Henry, D. Ji, A. Chambers, J. Warren, W. Shu, E.J. Weinstein, X. Cui, Whole-rat conditional gene knockout via genome editing, Nat Methods 10(7) (2013) 638–40.

[63] S.V. Mahler, Z.D. Brodnik, B.M. Cox, W.C. Buchta, B.S. Bentzley, J. Quintanilla, Z.A. Cope, E.C. Lin, M.D. Riedy, M.D. Scofield, J. Messinger, C.M. Ruiz, A.C. Riegel, R.A. Espana, G. Aston-Jones, Chemogenetic Manipulations of Ventral Tegmental Area Dopamine Neurons Reveal Multifaceted Roles in Cocaine Abuse, J Neurosci 39(3) (2019) 503–518.

[64] J.L. Gomez, J. Bonaventura, W. Lesniak, W.B. Mathews, P. Sysa-Shah, L.A. Rodriguez, R.J. Ellis, C.T. Richie, B.K. Harvey, R.F. Dannals, M.G. Pomper, A. Bonci, M. Michaelides, Chemogenetics revealed: DREADD occupancy and activation via converted clozapine, Science 357(6350) (2017) 503–507.

[65] H.L. Robinson, K.L. Nicholson, K.L. Shelton, P.J. Hamilton, M.L. Banks, Comparison of three DREADD agonists acting on Gq-DREADDs in the ventral tegmental area to alter locomotor activity in tyrosine hydroxylase:Cre male and female rats, Behav Brain Res 455 (2023) 114674.

[66] L. Boekhoudt, A. Omrani, M.C. Luijendijk, I.G. Wolterink-Donselaar, E.C. Wijbrans, G. van der Plasse, R.A. Adan, Chemogenetic activation of dopamine neurons in the ventral tegmental area, but not substantia nigra, induces hyperactivity in rats, Eur Neuropsychopharmacol 26(11) (2016) 1784–1793.

[67] M. Bortolato, R. Frau, G.N. Aru, M. Orru, G.L. Gessa, Baclofen reverses the reduction in prepulse inhibition of the acoustic startle response induced by dizocilpine, but not by apomorphine, Psychopharmacology (Berl) 171(3) (2004) 322–30.

[68] M. Bortolato, R. Frau, M. Orru, M. Collu, G. Mereu, M. Carta, F. Fadda, R. Stancampiano, Effects of tryptophan deficiency on prepulse inhibition of the acoustic startle in rats, Psychopharmacology (Berl) 198(2) (2008) 191–200.

[69] R. Frau, L.J. Mosher, V. Bini, G. Pillolla, R. Pes, P. Saba, S. Fanni, P. Devoto, M. Bortolato, The neurosteroidogenic enzyme 5alpha-reductase modulates the role of D1 dopamine receptors in rat sensorimotor gating, Psychoneuroendocrinology 63 (2016) 59–67.

[70] P. Wong, Y. Sze, C.C. Chang, J. Lee, X. Zhang, Pregnenolone sulfate normalizes schizophrenia-like behaviors in dopamine transporter knockout mice through the AKT/GSK3beta pathway, Transl Psychiatry 5 (2015) e528.

[71] S. Lovestone, R. Killick, M. Di Forti, R. Murray, Schizophrenia as a GSK-3 dysregulation disorder, Trends Neurosci 30(4) (2007) 142–9.

[72] J.M. Beaulieu, T.D. Sotnikova, W.D. Yao, L. Kockeritz, J.R. Woodgett, R.R. Gainetdinov, M.G. Caron, Lithium antagonizes dopamine-dependent behaviors mediated by an AKT/glycogen synthase kinase 3 signaling cascade, Proc Natl Acad Sci U S A 101(14) (2004) 5099–104.

[73] P.S. Klein, D.A. Melton, A molecular mechanism for the effect of lithium on development, Proc Natl Acad Sci U S A 93(16) (1996) 8455–9.

[74] K. Bolhuis, S.A. Kushner, S. Yalniz, M.H.J. Hillegers, V.W.V. Jaddoe, H. Tiemeier, H. El Marroun, Maternal and paternal cannabis use during pregnancy and the risk of psychotic-like experiences in the offspring, Schizophr Res 202 (2018) 322–327.

[75] D.J. Corsi, J. Donelle, E. Sucha, S. Hawken, H. Hsu, D. El-Chaar, L. Bisnaire, D. Fell, S.W. Wen, M. Walker, Maternal cannabis use in pregnancy and child neurodevelopmental outcomes, Nat Med 26(10) (2020) 1536–1540.

[76] S.E. Paul, A.S. Hatoum, J.D. Fine, E.C. Johnson, I. Hansen, N.R. Karcher, A.L. Moreau, E. Bondy, Y. Qu, E.B. Carter, C.E. Rogers, A. Agrawal, D.M. Barch, R. Bogdan, Associations Between Prenatal Cannabis Exposure and Childhood Outcomes: Results From the ABCD Study, JAMA Psychiatry 78(1) (2021) 64–76.

[77] D.C. Hoffman, H. Donovan, D1 and D2 dopamine receptor antagonists reverse prepulse inhibition deficits in an animal model of schizophrenia, Psychopharmacology (Berl) 115(4) (1994) 447–53.

[78] A.M. McCoy, T.D. Prevot, M.Y. Mian, J.M. Cook, A. Frazer, E.L. Sibille, F.R. Carreno, D.J. Lodge, Positive Allosteric Modulation of alpha5-GABAA Receptors Reverses Stress-Induced Alterations in Dopamine System Function and Prepulse Inhibition of Startle, Int J Neuropsychopharmacol 25(8) (2022) 688–698.

[79] N.R. Swerdlow, R.S. Mansbach, M.A. Geyer, L. Pulvirenti, G.F. Koob, D.L. Braff, Amphetamine disruption of prepulse inhibition of acoustic startle is reversed by depletion of mesolimbic dopamine, Psychopharmacology (Berl) 100(3) (1990) 413–6.

[80] B.B. Tournier, N. Ginovart, Repeated but not acute treatment with Δ(9)-tetrahydrocannabinol disrupts prepulse inhibition of the acoustic startle: reversal by the dopamine D(2)/(3) receptor antagonist haloperidol, Eur Neuropsychopharmacol 24(8) (2014) 1415–23.

[81] S.B. Caine, M.A. Geyer, N.R. Swerdlow, Effects of D3/D2 dopamine receptor agonists and antagonists on prepulse inhibition of acoustic startle in the rat, Neuropsychopharmacology 12(2) (1995) 139–45.

[82] N.R. Swerdlow, B.K. Lipska, D.R. Weinberger, D.L. Braff, G.E. Jaskiw, M.A. Geyer, Increased sensitivity to the sensorimotor gating-disruptive effects of apomorphine after lesions of medial prefrontal cortex or ventral hippocampus in adult rats, Psychopharmacology (Berl) 122(1) (1995) 27–34.

[83] S. Kapur, Psychosis as a state of aberrant salience: a framework linking biology, phenomenology, and pharmacology in schizophrenia, Am J Psychiatry 160(1) (2003) 13–23.

[84] O.D. Howes, R. McCutcheon, M.J. Owen, R.M. Murray, The Role of Genes, Stress, and Dopamine in the Development of Schizophrenia, Biol Psychiatry 81(1) (2017) 9–20.

[85] H. Sotoyama, H. Namba, Y. Kobayashi, T. Hasegawa, D. Watanabe, E. Nakatsukasa, K. Sakimura, T. Furuyashiki, H. Nawa, Resting-state dopaminergic cell firing in the ventral tegmental area negatively regulates affiliative social interactions in a developmental animal model of schizophrenia, Transl Psychiatry 11(1) (2021) 236.

[86] M. Kokkinou, E.E. Irvine, D.R. Bonsall, S. Natesan, L.A. Wells, M. Smith, J. Glegola, E.J. Paul, K. Tossell, M. Veronese, S. Khadayate, N. Dedic, S.C. Hopkins, M.A. Ungless, D.J. Withers, O.D. Howes, Reproducing the dopamine pathophysiology of schizophrenia and approaches to ameliorate it: a translational imaging study with ketamine, Mol Psychiatry 26(6) (2021) 2562–2576.

[87] C.S. Peterson, S.L. Baglot, N.A. Sallam, S. Mina, M.N. Hill, S.L. Borgland, Oral pre- and early postnatal cannabis exposure disinhibits ventral tegmental area dopamine neuron activity but does not influence cocaine preference in offspring in mice, J Neurosci Res 102(7) (2024) e25369.

[88] M.H. Sarikahya, S. Cousineau, M. De Felice, K. Lee, K.K. Wong, M.V. DeVuono, T. Jung, M. Rodriguez-Ruiz, T.H.J. Ng, D. Gummerson, E. Proud, D.B. Hardy, K.K. Yeung, W. Rushlow, S.R. Laviolette, Prenatal THC Exposure Induces Sex-Dependent Neuropsychiatric Endophenotypes in Offspring and Long-Term Disruptions in Fatty-Acid Signaling Pathways Directly in the Mesolimbic Circuitry, eNeuro 9(5) (2022).

[89] M. Ironside, P. Kumar, M. Kang, D. Pizzagalli, Brain mechanisms mediating effects of stress on reward sensitivity, Current Opinion in Behavioral Sciences 22 (2018) 106–113.

[90] D. Payer, B. Williams, E. Mansouri, S. Stevanovski, S. Nakajima, B. Le Foll, S. Kish, S. Houle, R. Mizrahi, S.R. George, T.P. George, I. Boileau, Corticotropin-releasing hormone and dopamine release in healthy individuals, Psychoneuroendocrinology 76 (2017) 192–196.

[91] A. Bara, J.N. Ferland, G. Rompala, H. Szutorisz, Y.L. Hurd, Cannabis and synaptic reprogramming of the developing brain, Nat Rev Neurosci 22(7) (2021) 423–438.

[92] R.J. Ellis, A. Bara, C.A. Vargas, A.L. Frick, E. Loh, J. Landry, T.O. Uzamere, J.E. Callens, Q. Martin, P. Rajarajan, K. Brennand, A. Ramakrishnan, L. Shen, H. Szutorisz, Y.L. Hurd, Prenatal Delta(9)-Tetrahydrocannabinol Exposure in Males Leads to Motivational Disturbances Related to Striatal Epigenetic Dysregulation, Biol Psychiatry 92(2) (2022) 127–138.

[93] Y.L. Hurd, J.N. Ferland, Y. Nomura, L.A. Hulvershorn, K.M. Gray, C. Thurstone, Cannabis Use and the Developing Brain: Highs and Lows, Front Young Minds 11 (2023).

[94] Y.L. Hurd, O.J. Manzoni, M.V. Pletnikov, F.S. Lee, S. Bhattacharyya, M. Melis, Cannabis and the Developing Brain: Insights into Its Long-Lasting Effects, J Neurosci 39(42) (2019) 8250–8258.

[95] H. Szutorisz, Y.L. Hurd, High times for cannabis: Epigenetic imprint and its legacy on brain and behavior, Neurosci Biobehav Rev 85 (2018) 93–101.

[96] A. Bara, A. Manduca, A. Bernabeu, M. Borsoi, M. Serviado, O. Lassalle, M.N. Murphy, J. Wager-Miller, K. Mackie, A.L. Pelissier-Alicot, V. Trezza, O.J. Manzoni, Sex-dependent effects of in utero cannabinoid exposure on cortical function, Elife 7 (2018).

[97] T. Black, S.L. Baccetto, I.L. Barnard, E. Finch, D.L. McElroy, F.V.L. Austin-Scott, Q. Greba, D. Michel, A. Zagzoog, J.G. Howland, R.B. Laprairie, Characterization of cannabinoid plasma concentration, maternal health, and cytokine levels in a rat model of prenatal Cannabis smoke exposure, Sci Rep 13(1) (2023) 21070.

[98] E. Murru, G. Carta, C. Manca, M. Verce, A. Everard, V. Serra, S. Aroni, M. Melis, S. Banni, Impact of Prenatal THC Exposure on Lipid Metabolism and Microbiota Composition in Rat Offspring., Helyion (2024).

[99] A.F. Scheyer, M. Melis, V. Trezza, O.J.J. Manzoni, Consequences of Perinatal Cannabis Exposure, Trends Neurosci 42(12) (2019) 871–884.

[100] G.I. Robinson, F. Ye, X. Lu, S.R. Laviolette, Q. Feng, Maternal Delta-9-Tetrahydrocannabinol Exposure Induces Abnormalities of the Developing Heart in Mice, Cannabis Cannabinoid Res 9(1) (2024) 121–133.

[101] H. Chang, M. Perkins, L. Novaes, F. Qian, T. Zhang, P. Neckel, S. Scherer, R. Ley, W. Han, I. de Araujo, Stress-sensitive neural circuits change the gut microbiome via duodenal glands, Cell (2024).

[102] E.R. de Kloet, M.L. Molendijk, Floating Rodents and Stress-Coping Neurobiology, Biol Psychiatry 90(4) (2021) e19–e21.

[103] J.E. Sutherland, L.C. Burian, J. Covault, L.H. Conti, The effect of restraint stress on prepulse inhibition and on corticotropin-releasing factor (CRF) and CRF receptor gene expression in Wistar-Kyoto and Brown Norway rats, Pharmacol Biochem Behav 97(2) (2010) 227–38.

[104] Y. Oda, N. Kanahara, H. Kimura, H. Watanabe, K. Hashimoto, M. Iyo, Genetic association between G protein-coupled receptor kinase 6/beta-arrestin 2 and dopamine supersensitivity psychosis in schizophrenia, Neuropsychiatr Dis Treat 11 (2015) 1845–51.

[105] M. Tamura, J. Mukai, J.A. Gordon, J.A. Gogos, Developmental Inhibition of Gsk3 Rescues Behavioral and Neurophysiological Deficits in a Mouse Model of Schizophrenia Predisposition, Neuron 89(5) (2016) 1100–9.

[106] L. Stertz, J. Di Re, G. Pei, G.R. Fries, E. Mendez, S. Li, L. Smith-Callahan, H. Raventos, J. Tipo, R. Cherukuru, Z. Zhao, Y. Liu, P. Jia, F. Laezza, C. Walss-Bass, Convergent genomic and pharmacological evidence of PI3K/GSK3 signaling alterations in neurons from schizophrenia patients, Neuropsychopharmacology 46(3) (2021) 673–682.

[107] S. Matsuda, Y. Ikeda, M. Murakami, Y. Nakagawa, A. Tsuji, Y. Kitagishi, Roles of PI3K/AKT/GSK3 Pathway Involved in Psychiatric Illnesses, Diseases 7(1) (2019).

[108] P. Wong, C.C. Chang, C.E. Marx, M.G. Caron, W.C. Wetsel, X. Zhang, Pregnenolone rescues schizophrenia-like behavior in dopamine transporter knockout mice, PLoS One 7(12) (2012) e51455.

[109] E.M. Hur, F.Q. Zhou, GSK3 signalling in neural development, Nat Rev Neurosci 11(8) (2010) 539–51.

[110] J.M. Beaulieu, A role for Akt and glycogen synthase kinase-3 as integrators of dopamine and serotonin neurotransmission in mental health, J Psychiatry Neurosci 37(1) (2012) 7–16.

[111] C.S. Karam, J.S. Ballon, N.M. Bivens, Z. Freyberg, R.R. Girgis, J.E. Lizardi-Ortiz, S. Markx, J.A. Lieberman, J.A. Javitch, Signaling pathways in schizophrenia: emerging targets and therapeutic strategies, Trends Pharmacol Sci 31(8) (2010) 381–90.

[112] P. Su, H. Zhang, A.H.C. Wong, F. Liu, The DISC1 R264Q variant increases affinity for the dopamine D2 receptor and increases GSK3 activity, Mol Brain 13(1) (2020) 87.

[113] Z. Freyberg, S.J. Ferrando, J.A. Javitch, Roles of the Akt/GSK-3 and Wnt signaling pathways in schizophrenia and antipsychotic drug action, Am J Psychiatry 167(4) (2010) 388–96.

[114] D.Z. Luo, C.Y. Chang, T.R. Huang, V. Studer, T.W. Wang, W.S. Lai, Lithium for schizophrenia: supporting evidence from a 12-year, nationwide health insurance database and from Akt1-deficient mouse and cellular models, Sci Rep 10(1) (2020) 647.

[115] P. Duda, J. Wisniewski, T. Wojtowicz, O. Wojcicka, M. Jaskiewicz, D. Drulis-Fajdasz, D. Rakus, J.A. McCubrey, A. Gizak, Targeting GSK3 signaling as a potential therapy of neurodegenerative diseases and aging, Expert Opin Ther Targets 22(10) (2018) 833–848.

